# The transgenerational consequences of paternal social isolation and predation exposure in threespined sticklebacks

**DOI:** 10.1101/2024.01.16.575739

**Authors:** Jennifer Hellmann, Michaela Rogers

**Affiliations:** The Ohio State University

**Keywords:** *Gasterosteus aculeatus*, paternal effects, phenotypic plasticity, predation risk, social buffering, stress, intergenerational

## Abstract

1. Parents routinely encounter stress in the ecological environment that can affect offspring development (transgenerational plasticity: TGP); however, parents’ interactions with conspecifics may alter how parents respond to ecological stressors.
2. During social buffering, the presence of conspecifics can reduce the response to or increase the speed of recovery from a stressor. This may have cascading effects on offspring if conspecifics can mitigate parental responses to ecological stress in ways that blunt the transmission of stress-induced transgenerational effects.
3. Here, we simultaneously manipulated both paternal social isolation and experience with predation risk prior to fertilization in threespined stickleback (*Gasterosteus aculeatus*). We generated offspring via in-vitro fertilization to allow us to isolate paternal effects mediated via sperm alone (i.e., in the absence of paternal care). If social buffering mitigates TGP induced by paternal exposure to predation risk, then we expect the transgenerational effects of predation exposure to be weaker when a conspecific is present compared to when the father is isolated.
4. Offspring of predator-exposed fathers showed reduced anxiety-like behavior and tended to be captured faster by the predator. Fathers who were socially isolated also had offspring that were captured faster by a live predator, suggesting that paternal social isolation may have maladaptive effects on how offspring respond to ecological stressors. Despite additive effects of paternal social isolation and paternal predation risk, we found no evidence of an interaction between these paternal treatments, suggesting that the presence of a conspecific did not buffer fathers and/or offspring from the effects of predation risk.
5. Our results suggest that socially-induced stress is an important, yet underappreciated, mediator of TGP and can elicit transgenerational effects even in species that do not form permanent social groups. Future studies should therefore consider how the parental social environment can affect both within and trans-generational responses to ecological stressors.

## Introduction

Organisms encounter a wide variety of threats and stressors in their environment that can push individuals beyond their homeostatic limits and induce physiological and behavioral changes that allow them to survive those stressors (Romero, Dickens & Cyr 2009). It is well known that individuals differ drastically in how they respond to these environmental stressors: the same external stressor can induce strong responses in some individuals and weak responses in others (Koolhaas *et al*. 1999; Cockrem 2013). Although some of this variation can be attributed to traits of the organism itself (e.g., personality (Cockrem 2013), sex (DeVries, Glasper & Detillion 2003)), some of this variation is environmentally dependent. In particular, there is ample evidence that the social environment alters how animals cope with ecological stress, influencing both the perception of and response to ecological cues (DeVries, Glasper & Detillion 2003; Brandl, Pruessner & Farine 2022). In many cases, the presence of conspecifics can reduce physiological and behavioral responses to stress via social buffering, which occurs when the presence of a conspecific reduces the response to or increases the speed of recovery from an external stressor (Kikusui, Winslow & Mori 2006; Gilmour & Bard 2022). For example, socially isolated mice showed greater immune impairment and a higher basal corticosterone increase in response to a mild stress compared to group housed mice (Bartolomucci *et al*. 2003). Similarly, in humans, anxiety and negative emotions induced by a social stress test were attenuated when women had social support (McQuaid *et al*. 2016). This social buffering may be an adaptive response to minimize the transmission of stress among group members (Gilmour & Bard 2022), which can occur when a stress response system (e.g., HPA axis) in one individual is activated after interacting with another stressed individual (Brandl, Pruessner & Farine 2022). Further, social buffering can even increase the limits of the physical conditions organisms are willing to tolerate (Gilmour & Bard 2022), suggesting that sociality can mediate responses to ecological stressors.

While social buffering has been well studied in highly social mammals, there is mounting evidence that social buffering of stress also occurs in less social species, including solitary species and species with aggregations or schooling behavior (Gilmour & Bard 2022). For example, in lake sturgeon (*Acipenser fulvescens*), group-housed fish had significantly reduced endocrine and cellular stress markers after exposure to heat stress compared to isolated individuals (Yusishen *et al*. 2020). This social buffering effect can even occur in small groups of two individuals (Faustino, Tacão-Monteiro & Oliveira 2017; Schumann *et al*. 2023) or in response to mere visual cues of a conspecific (da Costa *et al*. 2004; Faustino, Tacão-Monteiro & Oliveira 2017), suggesting that these social buffering effects are robust and occur in a wide range of taxonomic groups with differing levels of sociality.

While social buffering effects can have consequences for the individuals that directly experience the stressor, social buffering in parents may also have consequences for offspring. Transgenerational plasticity (TGP) occurs when the experiences of a previous generation alter the phenotypes of future generations (Bell & Hellmann 2019). TGP can occur in response to a variety of ecological stressors, including predation risk (Tariel, Plénet & Luquet 2020; MacLeod *et al*. 2022), heat exposure (Donelson *et al*. 2016; Pessato, McKechnie & Mariette 2022), immune challenges (Beemelmanns & Roth 2016; Moore, Kaletsky & Murphy 2019), and food limitation (Bygren *et al*. 2014; Hagmayer *et al*. 2022). These ecological stressors induce changes in parental state, including the activation of the hypothalamic-pituitary-adrenal axis (HPA axis), that influence offspring development, such as stress responsiveness, growth, and behavior (Bell & Hellmann 2019; Potticary & Duckworth 2020). Given that the magnitude of the stress response in parents can be linked to the magnitude of the phenotypic changes in offspring (Sheriff, Krebs & Boonstra 2010), social buffering may affect both the strength and frequency with which parental experiences affect offspring traits. This is supported by a recent study in humans which found that the adverse health outcomes differentiating offspring of healthy and psychologically stressed mothers could be specifically linked to a lack of social support in psychologically stressed mothers (Walsh *et al*. 2019). These effects may arise because conspecifics provide a buffering effect that abates maternal response to external stress (e.g., brings glucocorticoids back to baseline quickly (Kikusui, Winslow & Mori 2006)), mitigating adverse outcomes for offspring (Nylen, O’Hara & Engeldinger 2013).

Although this previous work suggests the importance of the social environment in mediating transgenerational effects for mothers, we know little about the transgenerational effects of social isolation for fathers. Fathers have at least two mechanisms by which they can influence offspring non-genetically: epigenetic changes to sperm/seminal fluid and behavioral changes to parental care (Immler 2018). Paternal experiences can affect offspring phenotypes in magnitudes that are similar to maternal experiences (Akkerman *et al*. 2016; Beemelmanns & Roth 2016; Hellmann *et al*. 2020), although it is difficult to compare the strength of parental effects in most systems because the effects of paternal experiences often have different effects on offspring than maternal experiences (Bonduriansky & Head 2007; Bouwhuis, Vedder & Becker 2015; Bonduriansky, Runagall-McNaull & Crean 2016; Bell & Hellmann 2019; Gilad & Scharf 2019; Hellmann *et al*. 2020). This is likely because males and females differ in their susceptibility to stress (DeVries, Glasper & Detillion 2003) and because the mechanisms of inheritance are different (e.g., eggs/placenta versus sperm). Consequently, the transgenerational effects of paternal isolation are likely different from maternal isolation. Thus, understanding how the social environment mediates paternal effects can help us better predict when and how paternal experiences influence offspring phenotypes and if, for instance, social buffering during or after a stressor mitigates or reverses changes in parental state that induce transgenerational effects.

We sought to understand the ways that the social environment alters the transgenerational consequences of paternal stress in threespined stickleback *(Gasterosteus aculeatus).* There is abundant evidence for predator-induced paternal effects mediated via sperm (i.e., in the absence of paternal care) in stickleback, which alter offspring behavior, growth, gene expression, survival, and stress responsiveness (Hellmann *et al*. 2020; Hellmann, Carlson & Bell 2020; Chen *et al*. 2021; Hellmann, Carlson & Bell 2021; Afseth *et al*. 2022; Hellmann *et al*. 2024). There is also previous evidence of social buffering effects in stickleback: fish that are exposed to simulated predation risk while isolated had higher circulating levels of cortisol compared to fish in a group (Mommer & Bell 2013) and pairs of sticklebacks co-regulate their cortisol responses in stressful environments (Fürtbauer & Heistermann 2016). However, we have very little understanding as to how the social environment might modulate parental responses to ecological stress in ways that might have cascading effects on the adaptive nature and frequency of transgenerational plasticity.

We used a full factorial design to compare the transgenerational consequences of exposing fathers to predation risk while fathers were isolated or with another conspecific. Offspring were generated via in-vitro fertilization; because fathers had no behavioral interactions with offspring post-fertilization, this allowed us to isolate paternal effects mediated via sperm and/or seminal fluid. We chose to focus on sperm-mediated paternal effects rather than paternal care because, in care-mediated paternal effects, it can be difficult to isolate changes in offspring phenotypes that come directly from the father (transgenerational effects) versus changes that result from the early life environment (within-generational effects). Further, a previous study found that the presence or absence of paternal care did not alter the transgenerational effects mediated via sperm alone (Hellmann, Carlson & Bell 2021). We then measured offspring behavioral traits relevant to predation defense, including anti-anxiety behavior, shoaling behavior, and survival against a live predator. If social buffering mitigates TGP induced by paternal exposure to predation risk, then we expect the transgenerational effects of predation exposure to be weaker when a conspecific is present compared to when the father is isolated.

## Materials and Methods

### Housing information and F0 exposures

In June 2021, we collected adult three-spined stickleback *(Gasterosteus aculeatus)* from Putah Creek (CA, USA), a freshwater stream in northern California, and shipped them to the University of Dayton. We maintained the F0 generation on a summer photoperiod schedule (16 L: 8 D) at 20° ± 1°C and fed them ad libitum daily with a mix of frozen bloodworms (*Chironomus* spp.), brine shrimp (Artemia spp.) Mysis shrimp, and Cyclop-eeze. We obtained 10-15 cm long rainbow trout *(Oncorhynchus mykiss)* from Freshwater Farms of Ohio in June 2021. Putah Creek has piscivorous predators, including the prickly sculpin (*Cottus asper*); while rainbow trout are present in Putah Creek, they are not present in the isolated section of Putah Creek where we collected stickleback (Beaver Pond), suggesting that rainbow trout is an environmentally relevant, but novel predator.

In August 2021, we moved males into experimental 37.9 L (53L x 33W x 25H cm) tanks divided into three equal sections using removable barriers. Males in the neighbor treatment were separated with clear barriers with nine 3mm holes to allow for water flow between the sections.

Males in the isolated treatment were separated with permanent opaque barriers that did not allow for water flow between sections. In both treatments, we transferred males from their home tank using a cup (males were never removed from water to minimize stress) and placed one male stickleback in each of the outer sections and left the middle section empty. Each male section contained gravel, one fake plant, and nesting materials: a container of sand, a terra cotta pot for cover, and algae. The middle section contained only gravel. Males that shared a tank were assigned to the same treatment group. Next to each nesting male tank, we placed an undivided 37.9L tank containing gravel and two sponge filters, which either contained one live rainbow trout or was left empty depending on the predator treatment. All experimental tanks were separated visually from each other with a removable opaque divider.

We exposed male stickleback to predator cues once a day for two hours, for 11-14 consecutive days depending on the availability of gravid females. We used a short stressor to avoid potential habituation to predation risk (Dellinger *et al*. 2018). Further, by beginning the treatment when males were transferred to a nesting arena, we sought to mimic the change in predator regime that males may encounter as they move into a different habitat to nest. At a random time of day between 0800 hrs and 2000 hrs, we turned off the bubblers in the tanks and removed the opaque divider to reveal the adjacent tank (which was empty or contained a live trout). At the same time, we added either 100mL of control RO water or the prepared predator cue to each end compartment containing one male stickleback. We prepared predator cues using methods consistent with Crane and Ferrari (2015). Briefly, we placed a trout in a 37.9 L tank with clean, conditioned tap water proportional to the size of the trout (500mL per gram of fish). We fed the trout one juvenile stickleback per day for four days and after 96hrs, we froze the tank water into 200 mL aliquots.

In the predator-naïve treatment, males saw an empty tank and received 100 mL of RO water. In the predator treatment, males saw a live trout and received 100 mL of the prepared predator cue. Every other day, we fed the trout one juvenile stickleback during the two hour exposure to reinforce the visual cue. Importantly, despite holes between the sections, we found very limited diffusion of water between the sections (tested using dyed water) when bubblers were turned off, suggesting that these olfactory cues did not disperse throughout the tank during the predation exposure and therefore, the neighbor treatment did not receive weaker olfactory cues or predation risk than the isolated group. However, water did disperse between sections when the bubblers were on, suggesting that neighboring males exchanged olfactory cues outside of the two-hour exposure period. After two hours, we replaced the dividers and completed a 50% water change on all treatments. We removed males after the last day of exposure.

### Generating Offspring

The day after the last exposure, we generated offspring by manually squeezing the eggs out of a gravid female (n = 29 different females across the 35 clutches, each sired by a unique male) and fertilizing them with sperm from the testes of the dissected experimental male. Females were all predator naïve and housed in group tanks on recirculating racks with biological, particulate, and UV filters. We checked clutches for successful fertilization via microscope and then placed into a small cup with a mesh bottom above an air bubbler in an 18.9L (38L x 25W x 21H) tank containing gravel and two fake plants, with each clutch housed individually on a recirculating system. By artificially fertilizing the eggs and incubating the embryos using an air bubbler, we controlled for possible pre-fertilization effects mediated by interactions between mothers and fathers (Mashoodh *et al*. 2012; McGhee *et al*. 2015), post-fertilization effects mediated by paternal care (Stein & Bell 2014), and the possibility that stressed parents might be less likely to successfully mate or parent offspring. Once hatched, we fed offspring newly hatched brine shrimp for two months and then gradually transitioned to frozen food (as above). We switched offspring to a winter light schedule (8 L: 16 D) prior to starting behavioral assays. We generated a total of n = 35 clutches: n = 9 of predator-naïve fathers with a neighbor, n = 9 of predator-naïve, isolated fathers, n = 8 of predator-exposed fathers with a neighbor, and n = 9 of predator-exposed, isolated fathers.. We assayed offspring at approximately 5 months of age; all offspring went through the shoaling assay first, followed by the scototaxis assay 24 hours later, and then the survival assay 48-72 after that.

### Offspring assays

#### Shoaling assays

We used experimental 37.9L (53 x 33 x 24 cm high) tanks lined with gravel. On one end of the tank, we positioned three small plastic plants; on the other end of the tank, we created a small compartment for shoaling fish by inserting a clear plastic barrier (10 cm from the edge) perforated with small holes to allow for flow of water and olfactory cues. The evening before a day of trials, we placed five size-matched juveniles (similarly aged juveniles that were not part of the experiment) into the shoaling side compartment of each tank to habituate overnight. These fish were unrelated and unfamiliar to the focal fish.

At the start of the shoaling trial, we placed an experimental offspring (focal fish) into the open area in the middle of the tank and allowed the focal fish to habituate for 10 minutes. After the habitation, we used BORIS (Friard & Gamba 2016) to record total time spent shoaling, total time spent oriented to the shoal while shoaling (the focal fish’s head had to be pointing in the direction of at least one member of the shoal), total time spent hiding in plants, and time spent in the gravel in the center of the tank over a 10 minute period. We defined a focal fish to be shoaling or hiding in the plant if it was within one body length of the shoal barrier or plants, respectively. Upon completion of the assay, we gently removed the focal fish from the tank with a cup, weighed and measured them, and then placed them into individual holding tanks. We assayed and measured n = 242 fish: n = 61 of predator-naïve fathers with a neighbor, n = 57 of predator-naïve, isolated fathers, n = 66 of predator-exposed fathers with a neighbor, and n = 58 of predator-exposed, isolated fathers.

#### Measuring anxiety/cautiousness

Scototaxis (light/dark preference) protocols have been developed to test anti-anxiety/cautious behavior in fish (Maximino *et al*. 2010). On the day after the shoaling assay, offspring were gently caught with a cup from their holding tank and placed in a clear cylinder (10.5cm diameter) in the center of a half-black, half-white tank (51L x 28W x 19H cm). After a five minute acclimation period, we lifted the cylinder, and fish explored the tank for 15 minutes, during which we measured the latency to enter the white section, total time in the white section, and thigmotaxis behavior (time spent within a body length of the wall) within the white section. We only measured thigmotaxis in the white area because we felt that thigmotaxis in a light, open area is more reflective of anxiety like behavior compared to thigmotaxis in a darker area. After the 15-minute testing period, we removed the fish from the tank and placed them back in their holding tank. We assayed n = 234 fish: n = 58 of predator-naïve fathers with a neighbor, n = 55 of predator-naïve, isolated fathers, n = 64 of predator-exposed fathers with a neighbor, and n = 57 of predator-exposed, isolated fathers.

#### Survival assay

48-72 hours after the scototaxis assay, we conducted survival assays in groups of two siblings (same paternal treatment and genetic family). We filled circular portable pools with conditioned tap water (1m diameter, 13 cm water depth) and evenly placed six plants around each arena. We placed an individually marked rainbow trout (n = 4) in each pool 24-hours before the first day of assays. We fasted the trout for 48 hours prior to being placed in the pools and we fed them one non-experimental juvenile stickleback after being placed in the pool to keep hunger levels standardized across days. On each assay day, we used each trout for two assays, once in the morning and once in the afternoon. For each assay, we placed two siblings in a clear cylinder (20.3 cm diameter) in the middle of the arena. After acclimating for 20 minutes, we lifted the cylinder and used a stopwatch to time how long it took the trout to capture one of the fish (survival time). We had a total of n=81 trials and we analyzed all trials where one of the fish was captured within one hour (n=79 trials: n = 26 of predator-naïve fathers with a neighbor, n = 13 of predator-naïve, isolated fathers, n = 22 of predator-exposed fathers with a neighbor, and n = 18 of predator-exposed, isolated fathers)

#### Statistical methods

To get an estimate of shoaling behavior, we used principal components analysis (R package factoextra (Kassambara & Mundt 2017)) to combine several behaviors: time spent hidden in the plant, time spent shoaling, time spent oriented to the shoal, and time spent in the gravel in the center of the tank. We extracted an eigenvalue of 2.74 that captured 68.5% of the variance in these behaviors; positive values indicate more social individuals who spent more time shoaling (contribution: 33.4%) and oriented to the shoal (31.3%), and spent less time hiding in plants (25.1%) and gravel (10.2%). This PC was used as the dependent variable in the model described below (Gaussian distribution). For the scototaxis assay, latency to enter the white arena was negatively correlated with time spent in the white section (Spearman rank correlation: ρ = – 0.42, p<0.001), so we analyzed variation in time spent in the white section of the tank, as well as time spent performing thigmotaxis in the white section of the tank (both Poisson distribution).

We used the package MCMCglmm (Hadfield 2010) to analyze offspring behavior, offspring length, and offspring mass. We ran models with a weak prior on the variance (V=1, nu=0.002) because our data were heteroskedastic. We ran models for 200,000 iterations, with a burn-in of 3000 iterations and thin = 3; we confirmed model fit and convergence. Models testing variation in shoaling and scototaxis behavior included fixed effects of paternal predation exposure, paternal isolation, and offspring standard length as well as random effects of clutch identity and observer identity. For the thigmotaxis model, we also included a fixed effect of total time in the white section of the tank. The survival model (time to capture, log-transformed, Gaussian distribution) included fixed effects of paternal predation exposure and paternal isolation, and random effects of clutch identity, observer identity (n = 4), and trout identity (n = 4). The model testing predictors of offspring length (Gaussian distribution) included fixed effects of paternal predation exposure, paternal isolation, and offspring age, while the model testing predictors of offspring mass (Gaussian distribution) included fixed effects of paternal predation exposure, paternal isolation, and offspring length; both models included random effects of clutch identity and observer identity. For all models, we included the interaction between paternal predation exposure and paternal isolation as this was part of our *a priori* predictions. We estimated effect sizes for all significant effects of paternal treatment using rank-epsilon-squared tests (R package effectsize (Ben-Shachar, Lüdecke & Makowski 2020)) to account for the non-parametric nature of our data. Effect sizes less than 0.06 indicate a small effect, while effect sizes between 0.06 and 0.14 indicate a medium effect size (Ben-Shachar, Lüdecke & Makowski 2020).

#### Animal welfare note

All methods were approved by Institutional Animal Care and Use Committee of University of Dayton (protocol ID 021-05), including the use of live predators.

## Results

We examined how paternal exposure to predation risk and paternal social isolation altered offspring behavior (shoaling and anti-anxiety behavior), size, and survival against live predators.

### Offspring behavior and size

We found no change in offspring shoaling behavior based on either paternal isolation or paternal predation exposure, although longer offspring spent more time shoaling than smaller offspring (Table 1). However, in the scototaxis assay testing anti-anxiety behavior, we found that offspring of predator-exposed fathers spent more time in the white section of the tank compared to offspring of predator-naïve fathers (Table 1; Figure 1; rank ε² = 0.03). We found no significant effect of paternal isolation on time spent in the white section (Table 1; Figure 1). We found no significant effect of paternal isolation or paternal predation exposure on offspring thigmotaxis behavior (Table 1). We also found no differences in offspring length or mass based on either paternal isolation or paternal predation exposure (Table 1).

**Figure 1:**
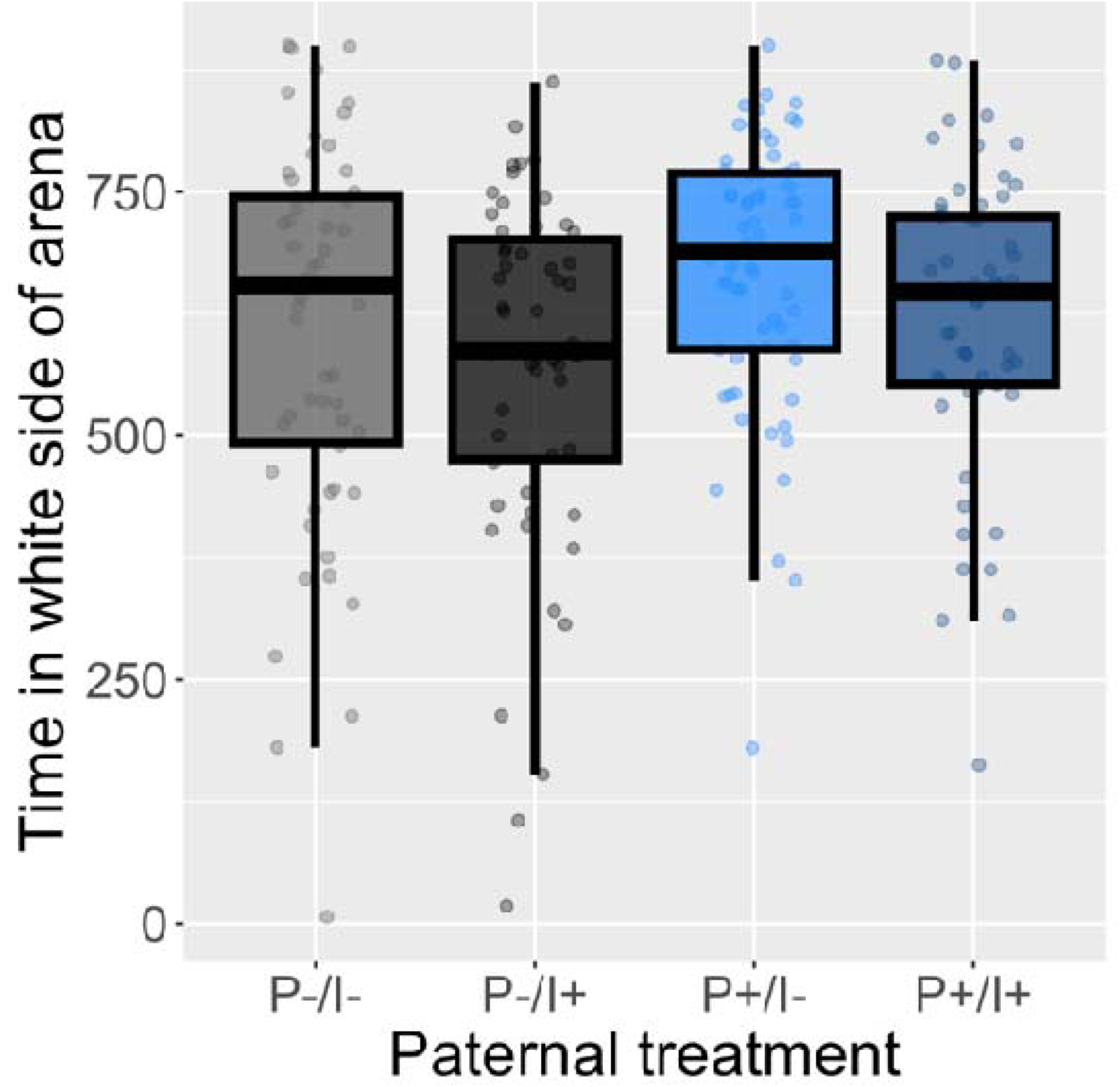
Offspring of predator-exposed fathers (blue) spent more time in the white section of the tank compared to offspring of predator-naïve fathers (grey), suggesting an anti-anxiety phenotype (median with interquartile range, dots correspond to individuals). P-indicates predator-naïve, unexposed fathers while P+ indicates predator-exposed fathers; I-indicates fathers who had a male neighbor, while I+ indicates isolated fathers.

**Table 1:** Results of mixed effects models on shoaling behavior (PCA), anti-anxiety behavior in the scototaxis assay (time in white section, thigmotaxis), survival against a live predator, mass, and length.

|  | **Shoaling** |  |  |
| --- | --- | --- | --- |
|  | *Mean* | *95% CI* | *P* |
| Paternal predation exposure | -0.10 | -0.80, 0.56 | 0.77 |
| Paternal isolation | -0.04 | -0.73, 0.61 | 0.89 |
| Standard length | 0.15 | 0.06, 0.24 | **0.001** |
| Paternal isolation * predation | -0.05 | -1.00, 0.91 | 0.92 |
|  | **Time in white section** |  |  |
|  | *Mean* | *95% CI* | *P* |
| Paternal predation exposure | 0.18 | 0.01, 0.36 | **0.047** |
| Paternal isolation | -0.05 | -0.24, 0.14 | 0.59 |
| Standard length | -0.01 | -0.03, 0.01 | 0.39 |
| Paternal isolation * predation | -0.04 | -0.29, 0.22 | 0.78 |
|  | **Thigmotaxis** |  |  |
|  | *Mean* | *95% CI* | *P* |
| Paternal predation exposure | 0.05 | -0.05, 0.16 | 0.33 |
| Paternal isolation | -0.01 | -0.13, 0.11 | 0.85 |
| Time in white section | 0.003 | 0.0028, 0.0033 | **<0.001** |
| Length | 0.02 | 0.005, 0.03 | **0.009** |
| Paternal isolation * predation | -0.01 | -0.17, 0.14 | 0.85 |
|  | **Survival time** |  |  |
|  | *Mean* | *95% CI* | *P* |
| Paternal predation exposure | -0.89 | -1.80, 0.07 | 0.058 |
| Paternal isolation | -1.17 | -2.24, -0.13 | **0.03** |
| Paternal isolation * predation | 0.64 | -0.85, 2.09 | 0.38 |
|  | **Length** |  |  |
|  | *Mean* | *95% CI* | *P* |
| Paternal predation exposure | 1.26 | -1.24, 3.74 | 0.31 |
| Paternal isolation | 0.04 | -2.50, 2.47 | 0.97 |
| Age | 0.05 | 0.01, 0.09 | **0.009** |
| Paternal isolation * predation | -1.31 | -4.81, 2.34 | 0.46 |
|  | **Mass** |  |  |
|  | *Mean* | *95% CI* | *P* |
| Paternal predation exposure | -0.005 | -0.02, 0.01 | 0.60 |
| Paternal isolation | -0.003 | -0.02, 0.01 | 0.72 |
| Length | 0.01 | 0.010, 0.0014 | **<0.001** |
| Paternal isolation * predation | 0.005 | -0.02, 0.03 | 0.71 |

### Offspring survival

We found that offspring of fathers who were isolated were captured significantly faster by the predator compared to offspring of fathers with neighbors (mean 45 sec ± 19 s.e. versus 153 sec ± 49 s.e.; rank ε² = 0.07; Table 1; Figure 2). Similarly, offspring of fathers who were predator-exposed tended to be captured significantly faster by the predator compared to offspring of predator-naïve fathers (mean 87 sec ± 49 s.e. versus 135 sec ± 39 s.e.; rank ε² = 0.07; Table 1; Figure 2). While the combined influence of paternal social isolation and predation risk seemed to have an additive effect on offspring survival (Figure 2), we did not find a significant interaction of paternal isolation and paternal predation exposure (Table 1), suggesting no evidence for social buffering.

**Figure 2.**
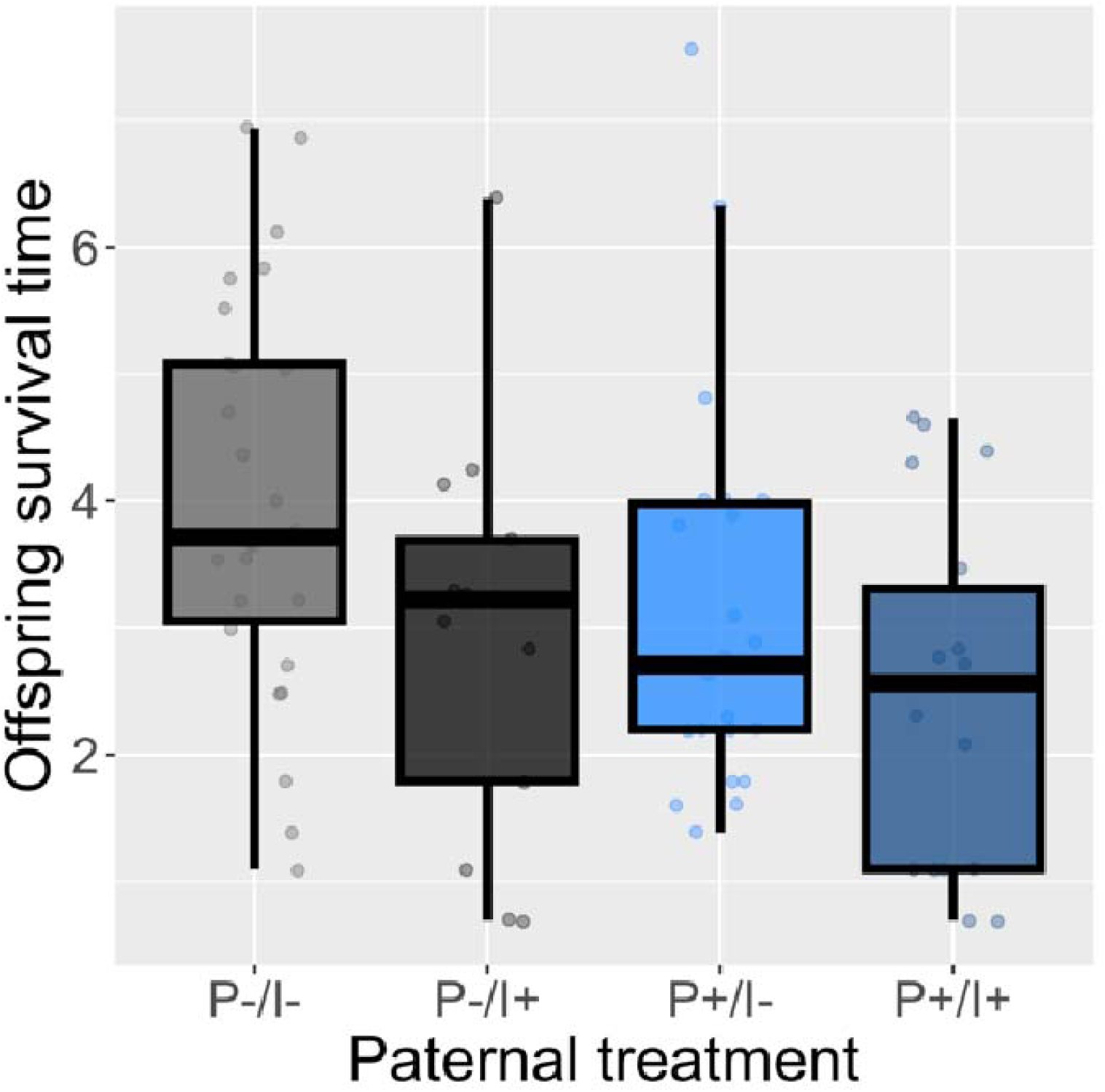
Offspring of socially isolated fathers (darker shade) were captured more quickly by the predator compared to offspring of a father with a male neighbor (lighter color). Similarly, offspring of predator-exposed fathers (blue) tended to be captured more quickly by the predator compared to offspring of predator-naïve fathers (grey). Data are median with interquartile range, dots correspond to individuals. Although raw integer data were used for the statistical analysis, we log-transformed the data here for better visibility of results.

## Discussion

Parents routinely encounter stress in the ecological environment that can affect offspring development. However, the social environment may alter how parents respond to ecological stressors, and therefore, affect how those parental experiences get integrated into offspring phenotypes. Here, we sought to understand how paternal social buffering (via the presence or absence of conspecific neighbors) altered the transgenerational consequences of paternal predation exposure. Offspring of predator-exposed fathers spent more time in the white section of the tank, consistent with an anti-anxiety phenotype, and had offspring that tended to be captured faster by the predator. We found that fathers who were socially isolated had offspring who were captured faster by a live predator, suggesting that social isolation in parents may be a stressor that alters offspring ability to cope with ecological stress. Despite evidence that paternal social isolation and predation risk have additive effects on offspring survival, we did not find evidence for a social buffering effect, where the presence of a conspecific buffers fathers and/or offspring from the effects of predation risk.

Previous studies have found that social isolation itself can have a detrimental effect on those directly experiencing it, including an immunosuppressive effect in greylag geese (Ludwig *et al*. 2017), oxidative stress in fish (Schumann *et al*. 2023), and elevated adrenocorticotropic hormone and corticosterone in mice (Dronjak *et al*. 2004). This likely occurs because social isolation can elevate glucocorticoids, which can have detrimental effects if individuals are chronically stressed (DeVries, Glasper & Detillion 2003). Here, we demonstrate that social isolation in adult males does not just affect individuals within a lifetime, but also has detrimental consequences for offspring, leading to lower survival against a predator. The effect size of paternal isolation on offspring survival was moderate and biologically meaningful, with offspring captured, on average, over three times faster when their father was isolated compared to when he had a neighbor. These maladaptive effects are consistent with a previous study in rats, where offspring of socially-isolated mothers showed cognitive impairments and increased anxiety-like behaviors (McDonald *et al*. 2023). These transgenerational effects of social isolation may arise because parental stress can hamper the ability of offspring to respond appropriately to stress encountered within their own lifetime (Yehuda *et al*. 2008; Rodgers *et al*. 2013; Faraji *et al*. 2017; McDonald *et al*. 2023), thus potentially leading to maladaptive behaviors in stressful situations (e.g., predation events (Hellmann *et al*. 2020)). Although no studies (to our knowledge) have looked at the transgenerational consequences of paternal isolation prior to fertilization, our results are consistent with previous studies demonstrating that other forms of paternal social stress, including maternal separation early in life (Gapp *et al*. 2014), social defeat (Dietz *et al*. 2011), and social instability (Kong *et al*. 2021), alter offspring anxiety and corticosterone levels. Consequently, although most TGP literature focuses on the effects of parental exposure to ecological stress (e.g., heat, predation, toxins, disease), we suggest that social stress can also have transgenerational consequences for offspring in ways that have not yet been fully appreciated.

The mechanism by which paternal social isolation affects offspring is unknown, but is clearly different than mechanisms of maternal social isolation, likely because those maternal social effects seem to be passed down via glucocorticoids in eggs or during gestation (Walsh *et al*. 2019; MacLeod *et al*. 2023; McDonald *et al*. 2023). Dietz *et al*. (2011) found that paternal social defeat in mice increased offspring susceptibility to depression and anxiety-like behaviors, but that this effect was largely not present when offspring were generated via IVF, suggesting that these transgenerational effects were mediated by changes in maternal investment in her offspring in response to perceived paternal quality (acquired during behavioral interactions between the mother and father). Here, we did not allow for the effects of maternal compensation on paternal phenotypes, as offspring were generated via in-vitro fertilization without behavioral interactions between the mother and father. Further, as our fathers also did not interact behaviorally with their offspring (no paternal care), our results demonstrate that these effects can be mediated via epigenetic changes to sperm and/or seminal fluid alone. Future studies linking the precise mechanism (e.g., small RNAs or methylation in sperm (Immler 2018)) to phenotypic effects in offspring would be useful for predicting when and how paternal social experiences affect offspring.

We found that offspring of predator-exposed fathers spent more time in the white side of the scototaxis arena, suggesting that they were less anxious (Maximino *et al*. 2010). This is consistent with previous studies demonstrating that paternal stress results in offspring with reduced anxiety and HPA responsiveness (Rodgers *et al*. 2013; Gapp *et al*. 2014; He *et al*. 2016; Brass *et al*. 2020), although results on this are mixed (Batchelor & Pang 2019). Further, offspring of predator-exposed fathers tended to be consumed faster by the predator, which is consistent with previous work in sticklebacks that found that paternal exposure to predation risk lowered offspring survival against a predator (Hellmann *et al*. 2020). While the effects of paternal isolation and predation exposure appeared to be additive, we found no evidence that the presence of a conspecific buffered fathers against the effects of predation risk (i.e., no evidence of an interaction between paternal predation exposure and paternal isolation). This was surprising, given abundant evidence that isolated individuals show high levels of stress in response to environmental stimuli, while individuals in social groups recover more quickly from or have a reduced response to stressful experiences (Dronjak *et al*. 2004; Kikusui, Winslow & Mori 2006; Ozbay *et al*. 2007; Culbert, Gilmour & Balshine 2019). Additionally, while some species do not show signs of social buffering (Gilmour & Bard 2022), there is previous evidence of social buffering in sticklebacks (Mommer & Bell 2013).

There may be several reasons why we did not see social buffering effects in this experiment. First, it is possible that the presence of a conspecific did alter how fathers responded to the presence of the predator, but not in a way that had consequences for offspring. Future studies linking paternal physiology and behavior to phenotypic effects in offspring would be useful for assessing this possibility. Second, it is possible that allowing for paternal care would have enhanced the social buffering effect, as the presence of a caring parent can buffer offspring against stressors (Sanchez, McCormack & Howell 2015). Finally, it is possible that the potential for social buffering was limited in our experiment, given that we paired two unfamiliar males together, both of whom were exposed to the predator. This would typically lead to weak effects of social buffering, as previous studies have found stronger effects of social buffering when individuals are paired with a known conspecific compared to an unknown conspecific (Hennessy, Kaiser & Sachser 2009), when individuals are in groups compared to pairs (Stanton, Patterson & Levine 1985; Kikusui, Winslow & Mori 2006), and when the partner is unstressed compared to stressed (Kiyokawa *et al*. 2004; Kikusui, Winslow & Mori 2006). Consequently, future studies manipulating the quality of the social environment could help us better understand how transgenerational effects of social buffering may vary with the number, familiarity, and state of the other conspecifics.

## Conclusions

We have long known that the social environment is an important source of selection, and that any given trait is not necessarily the property of a single individual because social interactions will influence the traits and fitness of other individuals, even if those individuals have never physically met (Fisher & McAdam 2017; Fisher *et al*. 2019). Currently, we view transgenerational plasticity as a linear and relatively isolated relationship between parent and offspring. Here, we demonstrate that the broader social environment can induce transgenerational effects even in relatively asocial species such as stickleback, which do not form permanent social groups. Although we did not find social buffering effects, previous work has shown that the social environment can mediate parental response to ecological cues, providing an opportunity for parents to learn about ecological change in their environment in ways that have cascading effects on offspring (Afseth *et al*. 2022). Further, we demonstrate that social stress can combine with ecological stress to have additive effects on offspring phenotypes. Future studies exploring the within and transgenerational consequences of these socially-induced stress effects, either by themselves or in conjunction with ecological stress, can help us better predict when and to what extent individuals will be able to adaptively response to rapidly changing environments.

